# Gendered Fitness Interests: A Method Partitioning the Effects of Family Composition on Socio-Political Attitudes and Behaviors

**DOI:** 10.1101/847814

**Authors:** Robert C. Brooks, Khandis R. Blake

## Abstract

Whereas most people are biologically either male or female, their genetic interests are almost never aligned with just their own sex. Drawing on the evolutionary theory of inclusive fitness gained through relatives, we partition the effects of kin on fitness into those that derive from female versus male relatives. We argue that the balance of these female- and male-derived effects, which we call ‘Gendered Fitness Interests’ (GFI), might influence human behavior, especially the adoption of socio-political attitudes that have a gendered dimension. Our proposal builds on observations that individual socio-political views covary with the sex of their children or the numbers of relatives of each sex. Further, it is consistent with the relatively small average differences between women’s and men’s socio-political positions. We outline a method for partitioning GFI, and use simulation models to explore some of its properties. We then show that (1) the GFI component of women’s and men’s socio-political attitudes will converge, on average, with age. (2) The contributions of both descendent and non-descendent kin lead to considerable variation in GFI. (3) When men have longer average reproductive lifespans than women, GFI can show small male-biases. (4) Paternity uncertainty reduces the variation in GFI between individuals, and (5) Large family sizes are associated with more variation among individuals in GFI. Our proposal provides a framework for the study of the effects of kin on traits and attitudes with a gendered dimension. In this respect, it may prove generally useful in resolving the complex origins of gendered behavior.

## Introduction

Some of the most ideologically polarizing issues faced by any society include a sex difference, with the probability that an individual will hold a particular view, or the fervor with which it is held, differing between women and men (Eysenck 1975, Feather 1977, Sidanius and Ekehammar 1980, Ekehammar and Sidanius 1982, DeVaus and McAllister 1989, Manza and Brooks 1998, Eagly et al. 2004). Women, for example, are more likely than men to favor gender equitable labor practices and remuneration, express punitive views on intimate partner violence, and favor public goods spending (Twenge 1997, Manza and Brooks 1998, Eagly et al. 2004, Eagly and Diekman 2006, Calvo-Salguero et al. 2008, Donnelly et al. 2016, Bell et al. 2018). Women and men also diverge, on average, on socio-political issues less directly related to reproduction and bodily autonomy, including the punishment of crime, the treatment of outgroups, economic redistribution, public goods spending, and religiosity (Eysenck 1975, Feather 1977, Sidanius and Ekehammar 1980, Ekehammar and Sidanius 1982, Prokos et al. 2010, Lizotte 2016, 2017). These issues assort with those more closely tied to reproduction and gender roles to define the progressive-conservative political axis (Pratto et al. 1994, Sidanius et al. 1994).

Many of these issues in which sex differences pertain have historically been considered ‘women’s issues’. That is, they affect women and men differently, and conservative positions often favor men’s interests over women’s. Hence the push for social progress, or for gender equity, is seen by some as a ‘woman’s issue’, with the ‘men’s position’ mapping onto the conservative side of the socio-political axis. On these issues, women and men often diverge in voting behavior (Eysenck 1975, Feather 1977, Sidanius and Ekehammar 1980, Ekehammar and Sidanius 1982, DeVaus and McAllister 1989, Eagly et al. 2003). Yet, there are many women who favor ‘conservative’ positions, and many men who favour ‘progressive’ positions. Moreover, while sex differences on such issues have been extensively documented in several societies (Eysenck 1975, Feather 1977, Sidanius and Ekehammar 1980, Ekehammar and Sidanius 1982, Prokos et al. 2010, Lizotte 2016, 2017), they are often quite small (see, for example, Hyde 2005, Eagly and Diekman 2006, Lizotte 2015). This fact suggests that sex shapes only a small portion of observed variation relative to other sources of individual variation including but not limited to wealth, status, race, class, education, religiosity, intelligence, and health.

Here we describe and model a source of variation in gendered attitudes that generates variability both within and among sexes as a result of what we call ‘Gendered Fitness Interests’ (GFI). Humans invest time, money and other resources in their kin, and strategic differences in these investments can be driven by relatedness considerations (Alexander 1982, 1987, Betzig and Lombardo 1992, Gaulin et al. 1997, Alvard 2003, Jeon and Buss 2007, Lieberman et al. 2007, Perry and Daly 2017, Antfolk et al. 2018). Our proposal and models aim to provide a method for partitioning these genetic relatedness considerations and applying them to the origins of variation in socio-political attitudes. In particular, we build on the early ideas of Alexander (1982, 1987) concerning how moral paradoxes arise from conflicts of reproductive interests, and the work of Betzig and Lombardo (1992) who drew attention to the importance of the balance of female and male kin on attitudes concerning abortion. We suggest how GFI might be detected by careful experimental and analytic work, partitioned from the other effects that shape an individual’s sociopolitical attitudes, and used to generate theoretic predictions.

This proposal depends on the assumption that there exist ideological positions that favour the interests of women and girls over those of men and boys, and other ideological positions that have the opposite effect. Such sex differences in ‘interests’ can arise due to differences in socialization, in access to status and resources, and in social roles (Eagly et al. 2004, Eagly and Diekman 2006, Diekman and Schneider 2010). They could also arise due to sex differences in the optimal conditions for reproductive success and, ultimately, evolutionary fitness (Alexander 1982, 1987, Betzig and Lombardo 1992, Borgerhoff Mulder and Rauch 2009, Shackelford and Goetz 2012). Although social and evolutionary accounts of human behavior are often treated as competing alternatives, no appropriately disinterested reader could deny the evidence that erecting a dichotomy is both fallacious and unhelpful (see Pinker 2002, Berenbaum et al. 2011, Wood and Eagly 2012, Schmitt 2015, Schmitt et al. 2017). The proposal and model we present here will remain agnostic with respect to the developmental, social, and genetic processes that give rise to sex differences in ‘interests’, while holding the assumption that some differences in ‘interests’ exist.

### The transformative nature of parenthood and other forms of kinship

The sexes of a person’s children and other kin have been shown to shift a person’s attitudes along the socio-political spectrum (see early review by Lundberg 2005). Firms led by CEOs who have daughters have been shown to adopt more socially and environmentally progressive corporate policies (Cronqvist and Yu 2017). Such firms are also more likely to appoint female directors to their boards than are firms whose CEOs have no daughters (Dasgupta et al. 2018). Venture capital firms led by senior partners who have daughters are also more likely to hire female partners (Gompers and Wang 2017), with positive consequences for overall firm performance.

Such kin effects are by no means confined to the corporate world. Warner (1991) found that support for feminism was greater among parents of daughters than those of sons. Parents, particularly fathers, who have only daughters express stronger support for public policies that address gender equity, whereas those policies gained the least support from men who have only sons (Warner and Steel 1999). The sex of one’s children can also affect adherence to conservative gender roles (Downey et al. 1994), attitudes concerning affirmative action (Prokos et al. 2010), and social dominance orientations (Pratto et al. 1994, Sidanius et al. 1994). “Daughter effects” are found in the political sphere as well. Representatives in the U.S. Congress who have daughters vote more progressively on bills concerning reproductive rights, provisions for working families, and tax-free education (Washington 2008). In Great Britain, parents of daughters vote more for left-wing parties than do parents of sons (Oswald and Powdthavee 2010).

Findings such as these are often viewed as effects of the offspring “changing” their parents via parents’ ongoing observation of, and social interaction with, their children (Miller and Glass 1989, Warner and Steel 1999, Washington 2008). The sex of a child might also impact other forms of social interaction experienced by parents, including different sporting and cultural activities, social interaction surrounding schools, etcetera. One might expect adolescent or adult offspring to have particularly large effects on their parents, given the years of social interaction, both with the child and with others as an indirect result of the child involved. Yet many of the effects outlined here are observed in parents of infants and toddlers (Warner 1991, Warner and Steel 1999), where these various pathways have had relatively little time to become fully realized.

The sex of a new family member provides what economists call an ‘exogenous shock’, a putatively random imposition of a change that permits, over a replicated sample of such changes, powerful inferences of causality. Several recent studies apply this approach to show that the sex of a newborn or fetus is associated with behavioral change (Oswald and Powdthavee 2010, Shafer and Malhotra 2011, Pogrebna et al. 2018). From a longitudinal dataset on British voting patterns, Oswald and Powthavee (2010) showed that the birth of a daughter can *cause* parents to move toward the political left, whereas the arrival of a son can do the opposite. Similarly, analysis of data from the U.S.A. National Longitudinal Study of Youth found that having a daughter causes men to weaken their support for conservative (what they call ‘traditional’) gender roles, although the sex of a child had no such effect on women’s attitudes (Shafer and Malhotra 2011). The exogenous shock of finding out the sex of one’s child at birth, or even *in utero*, can extend beyond changing attitudes or voting intentions to entire syndromes of behavior. Parents who find out they are having (or have just had) a daughter become almost twice as risk-averse as those who are having or have had sons (Pogrebna et al. 2018).

Recently we showed that despite Tunisian men being more supportive of mandatory Islamic veiling of women, women with more sons were more supportive of veiling, and more likely to wear veils themselves than women with fewer sons (Blake et al. 2018). In that paper we suggest that what we call “Gendered Fitness Interests” (GFI) may be an important consideration in future studies that combine social, evolutionary and economic ideas to study gender and its manifestations. In the case of Tunisian mothers, those with sons adopted a more typically masculine position than those with daughters, which we argue favors a restricted female sexuality that serves the interests of sons over those of daughters-in-law (Blake et al. 2018).

Gendered Fitness Interests could extend beyond an individual’s children. In a paper that precedes most of the examples we cited above, Betzig and Lombardo (1992) showed that people’s attitudes to abortion vary with the number of female kin in the 15–50 age group “at risk” of unwanted pregnancy. As this number increases, so does the adoption of pro-choice policies. Yet as the number of reproductive age male kin increases, so does the adoption of pro-life policies. Their analysis extends the highly politicized issue of abortion from being a clash of male and female interests to one where the composition of an individual’s broader family might be germane to the position that person takes. It is this idea, and Betzig and Lombardo’s (1992) paper, that sparked our interest in Gendered Fitness Interests.

### Gendered Fitness Interests

While most individuals are either male or female, the study of GFI explores the notion that their evolutionary fitness interests are almost never entirely aligned with just their own sex. Hamilton (1964a, b) showed that evolutionary fitness comprises not only personal fitness through numbers of descendent kin, but also inclusive fitness through all genetic relatives. The effect of a relative on an individual’s fitness is moderated by genetic relatedness, *r*, the probability of the focal individual and the relative sharing a given allele by common descent. Parent-offspring dyads share exactly half their alleles by descent. Full sibling dyads share, on average, half of their alleles by descent. On average, half-siblings share one quarter of their genes by common descent, as do grandparents with their grandchildren, and aunts/uncles with their nephews/nieces. First cousins share, on average, one eighth of their genes and so forth. Accordingly, an individual’s inclusive fitness should, *ceteris paribus*, be influenced four times more strongly by a full sibling than it is by a full first cousin.

The notion of inclusive fitness can be extended to include a gendered dimension that accounts for the sex of the kin through which an individual’s inclusive fitness is likely to be gained. Over large numbers of kin, and generations of yet-to-be-born kin, an individual’s fitness is likely to come equally through males and females. But in the space of a generation, or even a few years, currently living members of one sex may have far greater effect on an individual’s fitness than the other sex. The extent to which an individual’s future fitness on this timescale is likely to come through male or female lines depends on (1) the individual’s own sex, (2) the sexes of their kin, and (3) the likely future reproductive success (i.e. residual reproductive values) of the individual and each of those kin.

Consider a hypothetical example, where two unrelated women each have a single child but no other close surviving kin. Each woman has, in the language of evolutionary life-history theory, the same residual reproductive value (RRV, her expected number of future offspring, Fisher 1930, Stearns 1992), here denoted *n*. If *N* is the average number of offspring a woman has in her lifetime, let us assume that both women are at the age where women have had, on average 0.8*N* offspring, and can expect to have an average of *n* = 0.2*N* more.

The existing child of woman *a* is a daughter, *a_1_*. Assume *a_1_* is on the threshold of adulthood, where the average girl is likely to have *N* offspring in the future. Her contribution to her mother’s inclusive fitness is *Nr*, and because she is a daughter, *r*=0.5. Thus, woman *a* can expect to have inclusive fitness of 0.2*N* + 0.5*N* = 0.7*N*. Moreover, note that all of woman *a*’s fitness due to individuals currently alive will come through females.

Woman *b* has a son, *b_1_* who is also on the threshold of adulthood and thus has the same expected offspring number, *N*. Woman *b*’s expected fitness is also 0.7*N*, but only 2/7ths through females (i.e., herself = 0.2*N*) and 5/7ths will come through existing males (i.e. her son *b_1_* = 0.5*N*). Thus, woman *b* has 2.5 times more of her future fitness interests in living males than females.

The same approach can be used for individuals with more living kin. The fitness effects, *k*, of all *y* kin of sex *x*, including the individual her/himself, can be summed as:

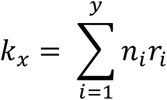

Where *n*_i_ is the expected future reproductive success of individual *i* as a proportion of the overall population mean reproductive success, and *r*_i_ is the relatedness of individual *i* to the focal individual. One might estimate *n_i_* by referring to the individual’s position on sex- and age-specific fecundity curves for the population they live in, and more complex treatments could include individual terms for mate value, survival and other traits.

One could then express the gendered nature of fitness interests either absolutely as a difference:

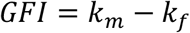

Now consider the simplified case of a focal individual with two children, and 100% paternity confidence, such that each child is related with *r* = 0.5 to the focal individual. Assume that every woman and man has 25 reproductive years (from age 20 to 44 inclusive), and that the likelihood of reproducing in any one of these years is the same. Thus *n* equals 1 at age 20, and declines monotonically to zero at age 45. Each child has 100 percent survival to reproductive age, so *n*=1 for all individuals younger than 20, and then from the age of 21 onwards, diminishes by 0.04 per year, until *n*=0 at age 45. Assuming no other living relatives, from birth until age 20 all of an individual’s future fitness will derive from their self. Thus Gendered Fitness Interests, expressed as *GFI*, equals -1 for women and 1 for men up to the age of 20.

Descendants begin to influence an individual’s GFI when they are born. In Figure 1 we illustrate how this simplified example might work for a man (Fig 1, Panel A) and a woman (Fig 1, Panel B), each of whom have one child at age 24 and a second at age 27, in relation to the sexes of those two children. The birth of each daughter reduces *GFI*, whereas the birth of a son raises *GFI*. Women and men with the same pattern of offspring have identical GFI after their own reproductive years end at 45 years.

**Figure 1.**
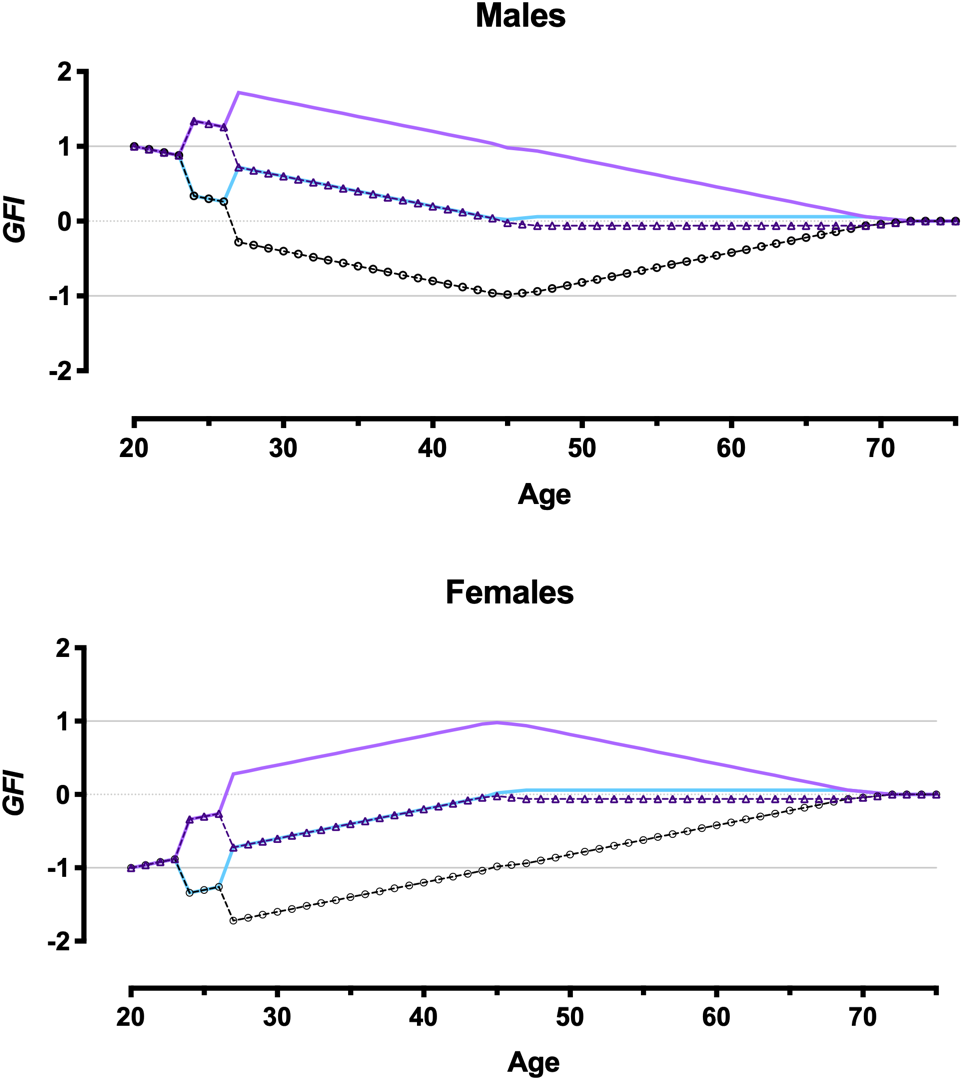
Examples of Gendered Fitness Interests for focal males (upper panel) and females (lower panel) in relation to age and the sex of two offspring born to them at ages 24 and 27. Dark purple solid line = son at 24, son at 27. Light purple dashed line and triangles = son at 24, daughter at 27. Light blue solid line = daughter at 24, son at 27. Black dashed line and circles = daughter at 24, daughter at 27.

As the focal individual’s offspring enter their reproductive years at 20, their own RRV starts declining and *GFI* begins moving toward zero. In reality most subjects will have grandchildren and other relatives, likely drawing their *GFI* away from zero but without the dramatic swerves of their own offspring due to lower relatedness.

We deliberately use an open-ended approach to quantifying GFI, in which individuals can and often do exceed the limits of -1 (a young female with no relatives) and 1 (a young male with no relatives). The effects of relatives are additive, such that, everything else being equal, a person with three daughters would have a far lower GFI until those daughters reach menopause, than a person who had only one daughter.

### Simulation: An Exploration of GFI (Model 1)

We built a simple simulation model in order to explore some of the properties of Gendered Fitness Interests for three generations, only including descendent kin (i.e., children and grandchildren) and assuming the focal individuals did not have living parents, siblings or other relatives. As in the previous example, we assume individuals can begin reproducing at age 20 and cease reproducing at 45. For simplicity, we divide the reproductive career into five-year intervals (20–24, 25–29 etc.). In each five-year interval, each individual could produce between 0 and 5 sons, by sampling at random from a Binomial distribution of 5 events, each with the probability, *p*, of a son being born as a result of that event. To do so we applied the BINOM.INV function in MS Excel with *Trials* = 5, *Probability_s* = *p*, and *Alpha* drawn as a random number (function *Rand()*). This is effectively the same as saying every focal individual could produce one son per year, at probability *p*. Independently, but otherwise in exactly the same way, each individual could produce between 0 and 5 daughters in each five-year interval.

From the interval 40–44 onward, offspring cohorts began to mature (i.e. reach 20 years old) and could produce grandchildren. We applied the same formula to allocating grandsons and, independently, granddaughters, but the number of trials was set to equal five times the number of offspring (both sons and daughters) of reproductive age. Thus each offspring of reproductive age could produce up to five grand offspring of each sex in a given 5-year interval.

We calculated *GFI* at the end of each five-year interval up to the age of 65. To simplify these calculations, we assume that offspring and grand-offspring are as old as it is possible to be in their cohort. For example, offspring born in the first cohort (parental age 20–24) are assumed to be 25 when the focal individual (i.e. parent) is 45. To confine our attention to three generations, we only present simulation results up to the focal individual age of 65. We present the outputs of simulations with *p* = 0.1, where each reproductive-age individual had a 10% chance of producing a son and an independent 10% chance of producing a daughter in a given year. This resulted in the following numbers (mean ± S.E; range) of descendent kin across a focal individual’s entire career: sons (2.49 ± 1.56; 0–5); daughters (2.52 ± 1.46; 0–5); grandsons (4.59 ± 3.47; 0–11); granddaughters (4.65 ± 3.54; 0–11).

The output of these simulations leads to a number of observations (see Figure 2a). First, GFI converges in males and females as the self’s residual reproductive value diminishes. By the time a focal individual reaches 45, reproduction is over and all fitness interests reside in descendants, male and female average gendered fitness interests are neutral (mean *GFI* = 0.5). Second, there remains considerable variation among individuals within sexes in GFI. The convergence of gendered fitness interests as we have conceptualized them leads to the prediction that attitudes that have a gendered dimension will show diminishing sex differences in mid- to late adulthood, and that they will retain a great deal of within-sex variation.

**Figure 2.**
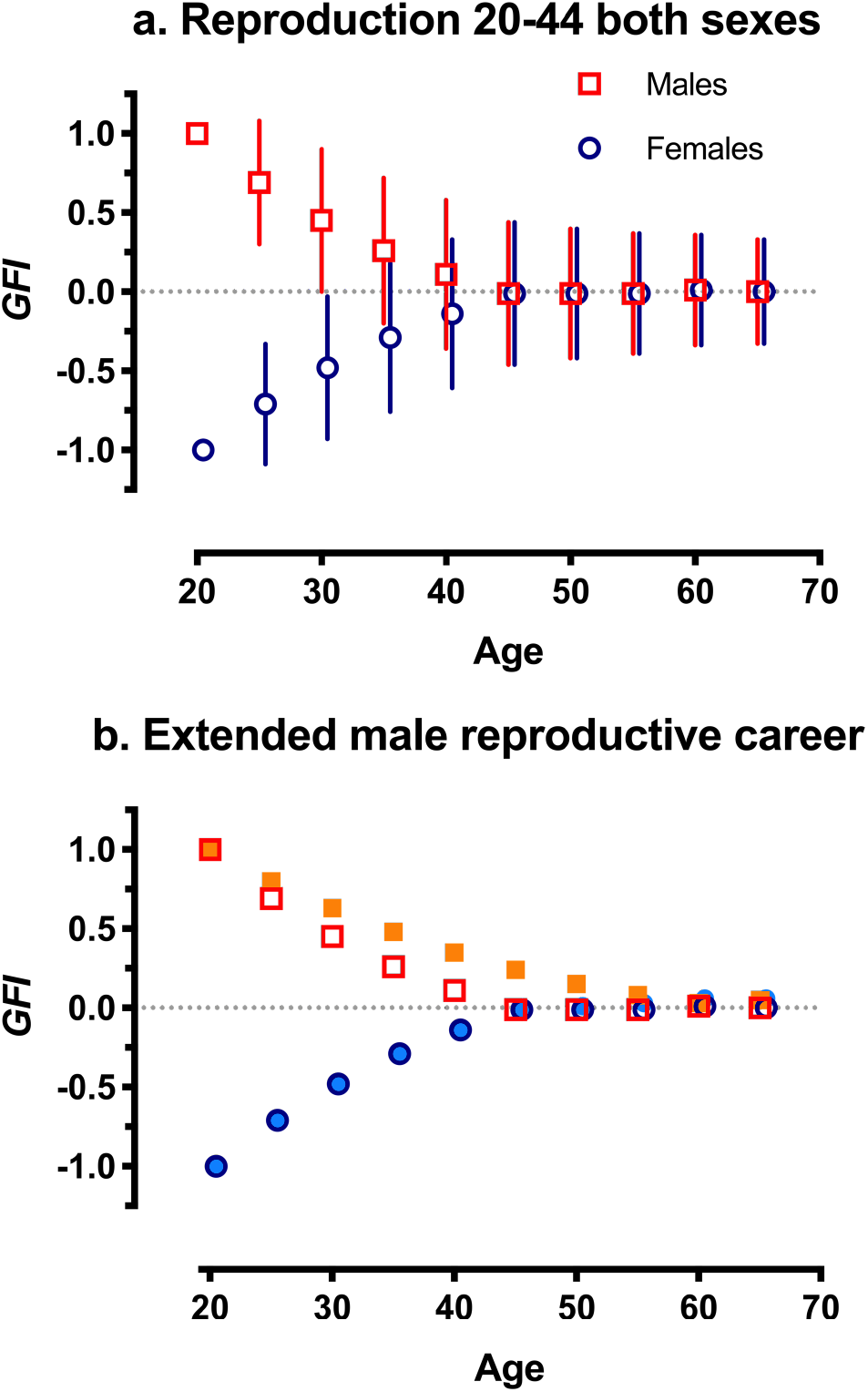
Output from 1000 runs of the three-generation descendent kin simulation models giving *GFI* at five-yearly intervals. A. Means and standard deviations for males (red squares) and females (dark blue circles) when both sexes can reproduce between ages 20-44 (Model 1). B. Means only for 25 year reproductive career (as in panel a), or for males (filled orange squares) and females (filled light blue circles) when males have a 40-year reproductive career (20-59 years old; Model 2). Females offset by 0.5 years; dotted line denotes equal fitness interests in males and females.

### Simulation with Unequal Career Durations (Model 2)

Our observed convergence at age 45 is an artificial result of our constraining reproduction to the 20–44 age range. Women often debut earlier than men as parents, and menopause curtails women’s reproduction from mid- to late adulthood whereas men peak later and can go on to reproduce at older ages than women. As a simple illustration of how different durations in men’s and women’s reproductive careers might alter gendered fitness interests, we altered the model above to allow men to reproduce for 15 years more than women (i.e., to 60 vs 45 years of age). The probability of a woman bearing a son in a given year of her 25-year career remained set at *p* = 0.1, and likewise the probability of her bearing a daughter in a given year equaled *p* = 0.1. For men, however, with a 40-year reproductive career these probabilities were scaled to *p* = 0.1 × (25/40) = 0.063, in order to satisfy the assumption that the population total reproductive output through males and females was the same. The results of 1000 simulations show that focal men were, on average, slower to converge toward *GFI* = 0.5 due to the longer persistence of the effects of the self (Figure 2b). Other modelling exercises (not presented) that further weight women’s reproduction to earlier ages and men’s reproduction to later years reveal even stronger biases toward positive (i.e. male-biased) GFI among descendent kin due to sons’ longer-lived contributions to GFI. The relationship between age- and sex-dependent fertility and GFI remains a fruitful avenue for future modelling.

### Simulation for Paternity Uncertainty (Model 3)

To examine how GFI is affected by male paternity uncertainty, we simulated GFI for women and men with offspring and grandoffspring. For each instance of paternity, the average relatedness of descendent kin to the focal individual was discounted by *u*, a probability between 0 and 1. A value of *u* = 0 indicates a father’s full paternity of the offspring, and *u*=1 indicates all offpsring are from misattributed paternity. Thus our simulations model the statistical paternity uncertainty, which accumulates over generations as the chance of at least one non-paternity event, rather than stochastically simulating individual paternities. As in previous models, we simulated reproduction by drawing from a binomial distribution. We drew the number of daughters, and, independently, the number of sons, assuming a 25-year reproductive career, an average final family size of 3, and only one offspring of each sex per year (i.e. the probability of producing an offspring of a given sex in a given year = 0.06). We ran 1000 simulations at each of the following values of *u*: 0, 0.05, 0.1. 0.15, 0.2, 0.5, 0.8 and 1. In the results (Fig 3), we convert these probabilities to percentages.

**Figure 3.**
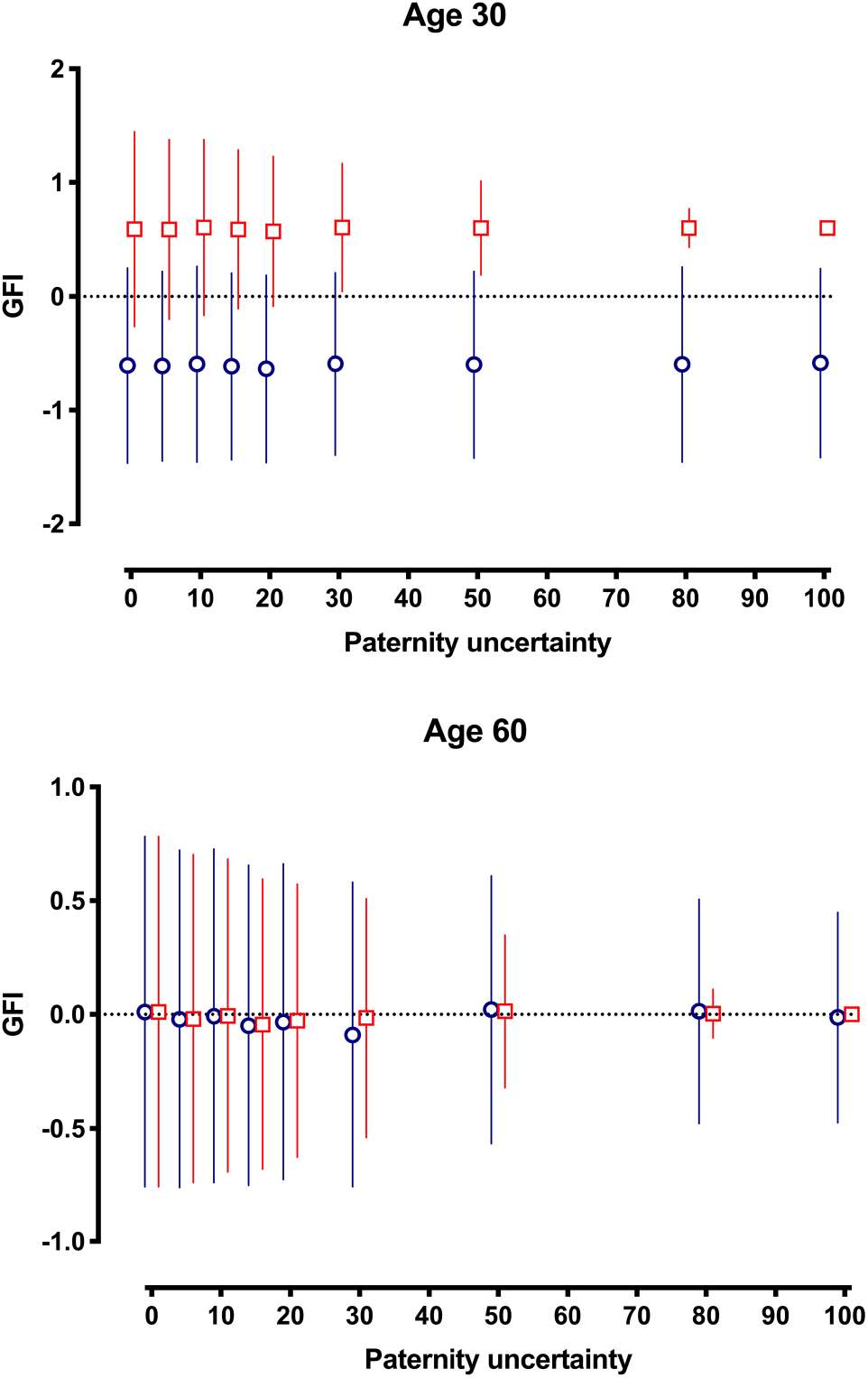
Gendered Fitness Interests in relation to varying levels of paternity uncertainty at ages 30 and 60. Output from 1000 runs of Model 3. Means for females (dark blue circles) and males (red squares) and Standard Deviations presented. At age 30, with only themselves and their offspring included, males have one opportunity and females no opportunities for misattributed paternity. At age 60, with grandchildren included, males have two opportunities and females one opportunity for misattributed paternity.

The results for focal individuals at age 30 and age 60 are presented in Figure 3. The mean differences between female and male focal individuals at age 30 are due to the effects of the self. At age 60 the effects of the self have declined to zero, and both male and female means tend toward GFI = 0. Paternity uncertainty had only trivial effects on mean GFI. It did, however, effect the variation about the mean for males at both ages and females at age 60, with greater paternity certainty (i.e., lower *u*) associated with more variation in GFI. This is because high paternity certainty ensures the full value of kin contributions to GFI, whereas lower paternity certainty discounts those contributions. It is the contributions of kin that are responsible for within-sex variation in GFI. Note that paternity certainty had no chance to effect female GFI at age 30 in this model because females had only their own offspring, and thus no opportunity for paternity misattribution. At age 60 females had one opportunity for paternity misattribution, through their sons. Males, by contrast, had one opportunity for paternity misattribution by age 30, and two by age 60. In effect, high levels of within-sex variation in GFI arise under lower levels of paternity uncertainty, particularly for males.

### Simulation for Non-descendent Kin and Family Sizes (Model 4)

Having simulated the effects of a few simple sources of variation on GFI, we sought to simulate GFI in a broader network of descendent and non-descendent kin, and, in the same simulation, to model the effects of family size on GFI. In this model we simulated families over five generations, starting with a female and male progenitor (P, or Parental generation), through four filial generations (F1 to F4). We track the GFI of P, F1 and F2 individuals of each sex at ages 20, 30, 40 and 60.

To simulate the P generation, we followed the same procedure described in the paternity uncertainty model above, assuming a 25 year reproductive career. Instead of fixing mean family size and thus the probability of having a child of a given sex in a given interval, however, we tested for family size = *s* at 2, 2.5, 3, 3.5, and 4. Thus the probability of an individual having an offspring of a given sex in a given year is given by *s*/50 (i.e. 25 years, 2 sexes). As all individuals in a given simulation descended from the parentals, these simulations merely required that we track the number of individuals at each level of descent.

For focal individuals in the F1 and subsequent generations, we calculated the GFI of the eldest male and female. The parents in every generation are assumed to be alive until the end of their reproductive lifespan, and thus cancel one another’s effects on GFI. In addition to descendent kin, our simulation introduces siblings (*r*=0.5), neice/nephews (0.125), and grand neice/nephews (0.0625) in the F1 generation, and uncles/aunts (0.25), cousins (0.125), and second cousins (0.0625) in the F2 generation.

The results in Figure 4 show the same pattern as in previous models, with the mean GFI of female and male focal individuals converging after the end of their own reproductive careers, but with considerable variation within each sex due to the effects of kin. One might have predicted that large families would experience fewer stochastic large GFI effects as large numbers of relatives cancel one another’s effects out. Instead, at larger mean family sizes, GFI was more variable than at smaller mean family sizes. This is likely due to an individual’s offspring and siblings having large effects on GFI due to their high relatedness, combined with the fact that extreme familes with large sex-biases (e.g., 5 daughters, no sons) are more likely, and families with no children are less likely at larger values of *s*.

**Figure 4.**
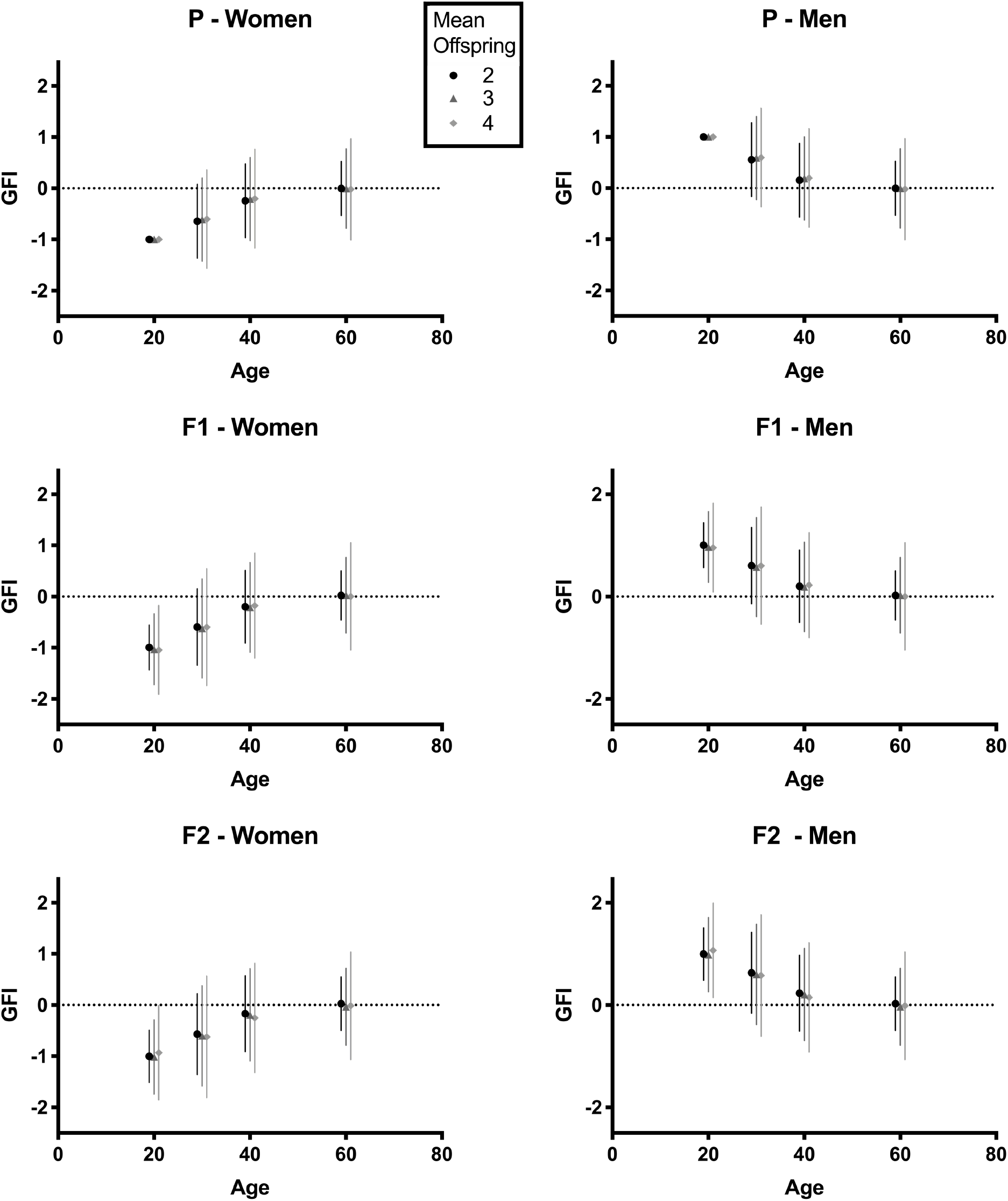
Effects of mean family size, sex, and age on GFI in a model incorporating non-descendent kin for three generations of focal individuals. Output from 1000 runs of Model 4. Means and standard deciations for families of average size 2 (Black circles), 3 (dark gray triangles) and 4 (light gray diamonds). Parental individuals have only themselves at age 20, and their descendent kin thereafter. F1 and F2 individuals have non-descendent kin.

Intriguingly, the distribution of GFI did not seem to differ greatly between the generations apart from the earliest measure of the parental generation in which GFI was determined solely by the sex of the self. This result suggests that non-descendent kin do not exert large leverage on GFI as we have operationalized it here.

Future attempts to model GFI will need to account for the complex effects of demographic factors, notably sex differences in age-specific fecundity. Differences in the age of debut, the age at which reproduction peaks, and the properties of its decline in each sex will alter the trajectory of GFI across an individual’s lifetime, as well as the average differences in GFI between women and men. The accurate estimation of GFI, especially as siblings and other non-descendent kin are also accounted for, is likely to become quite complex. It is our hope that theoreticians will develop more extensive models and predictions, and that empiricists will, in time, conduct direct tests of those predictions.

## Discussion

The examples above illustrate some simple ways in which the sex of an individual’s genetic kin might alter the inclusive fitness stake an individual has in females versus males. An individual’s own age (a proxy for residual reproductive value), the sex and age of any children they have, as well as the age and sex of their other relatives, will determine the overall strength of the GFI component of their attitudes. Here our simple modeling presents some preliminary attempts to understand how GFI might behave, and might be partitioned from other likely sources of variance in traits such as socio-political attitudes. We welcome more comprehensive attempts to understand GFI and how to tackle the complex job of estimating it, as well as to extract predictions for empirical testing.

We predict that individuals will track GFI, or convenient heuristics of GFI, and that their attitudes and other behaviors will change during the individual’s life as a consequence of changing GFI. This prediction remains largely untested (bust see Blake at al 2018). For now, our modeling gives rise to several observations and thus to some simple testable predictions.

### Sex Differences Diminish with Age

Our first observation is that an individual’s GFI changes as they get older, as they and their surviving kin age, and as other kin are born or die. This change in GFI will, on average, be toward the typical position for the other sex, thus narrowing the average difference between the sexes. This expectation is due both to a weakening in the effect of their self on GFI, due to diminishing residual reproductive value, and to the sex ratios of kin, all of which may cause GFI to regress toward the mean sex ratio (approximately 0.5). As a result, gaps in attitudes between women and men should, on average, be bigger in early adulthood and converge with age.

Despite a rich literature on the gendered nature of attitudes and orientations (e.g. Twenge 1997, Eagly et al. 2004, Calvo-Salguero et al. 2008), there is a surprising dearth of evidence on how individual attitudes change with age, and no evidence on the effects of kin on those changes. The majority of studies on attitudes have, for pragmatic reasons, been conducted on undergraduates or young adults (e.g., Sidanius and Ekehammar 1980, Ekehammar and Sidanius 1982, Twenge 1997). Those studies that have considered older adults, and the relationship between age and sociopolitical attitudes (e.g, Rice and Coates 1995, Cornelis et al. 2009), have usually involved crosssectional comparisons. Valuable as these cross-sectional data can be, they confound cohort effects with the within-individual change over time that is of interest to our hypothesis.

In one of the few studies to take a longitudinal view of attitude change over time, and also test for interactions between age and sex, Judge and Livingston (2008) found that people became less conservative in their gender role attitudes over time and that this effect was stronger for men. Thus, although women are, as expected, less conservative in their gender role attitudes than men, and there is an overall trend toward less conservative attitudes (likely due to secular change), men’s attitudes do appear to change faster than women’s, and the sex difference diminishes. This finding is consistent with our predictions.

Testing our prediction regarding age and attitude change will require longitudinal studies of individuals in order to separate true changes in attitudes from cohort effects. Convergence in attitudes is a prediction not unique to our proposed GFI mechanism. Women and men might be expected to converge due to all manner of shared experiences and socialization. Preferably, empirical tests will also track the sex and age of subjects’ close kin in order to separate effects of GFI from other sources of convergence. The perfect test, though laborious, will follow individuals longitudinally through a variety of exogenous changes in GFI due to births and unanticipated deaths, extending the approach used for single births by Oswald and Powthavee (2010), and Shafer and Malhotra (2011). Note that we make no claim that GFI is the only source of variance in sociopolitical attitudes, only that it could be important. Thus we predict a narrowing of the mean gap between the sexes with age as the effects of the self on *GFI* diminish. We do not necessarily predict complete closing of the gap as factors other than GFI are almost certainly always at play.

Our predictions that sex differences will diminish with age extend beyond population means to differences between spouses. Alexander (1987) noted that spouses share, as a result of their mutual offspring, similar avenues to fitness, and thus that their world views might converge due to shared reproductive interests. Observations from our similar model are consistent with Alexander’s conjecture. Spouses are well known to evince highly similar socio-political attitudes (Bouchard 2009, Alford et al. 2011). Most of that similarity, however, is already present early on in the relationship (Alford et al. 2011), and does not appear to increase with the duration of the relationship (Martin et al. 1986). Spousal similarity would appear to arise mostly from assortative dating and mating (Hatemi et al. 2010, Huber and Malhotra 2017), although it would be informative to test how changes in GFI through shared kin (e.g. children) enhance spousal similarity, and whether changes due to the births of nephews, nieces, etc. might open up some spousal differences. It may also be interesting to consider how cultural differences in courtship, matchmaking, and marriage customs that impact the degree of spousal similarity, might be modified as GFI within couples converges. The results of our Model 4, however, suggest that more distal non-descendent kin like nephews/nieces do not exert much leverage on an individual’s GFI due to low relatedness.

There exists great scope for longitudinal analyses, including data that track shifting individual roles (as per Eagly et al. 2004, Wood and Eagly 2012), and shifting constellations of close non-kin (e.g. stepchildren, adoptees) and affinal kin, in order to quantify the importance of GFI relative to other candidate hypotheses.

### Relation to Behavior Genetic Evidence

One intriguing study of socio-political attitudes and age comes from the behavior genetic study of twins (Eaves et al. 1997). The resemblance in “conservatism” between juvenile twins (younger than 20 years old) is no greater in monozygotic (r=1) than dizygotic (r=0.5) twins, suggesting that the resemblance is largely due to the social influence of the family. The correlation rises through the teens for both types of twins, reaching approximately 0.6-0.65 at age 20. After the age of 20, however, the resemblance between dizygotic twins drops away to about 0.4 at age 40, where it remains until age 75. The correlation in conservatism between monozygotic twins remains at or just above 0.6. The authors interpret this finding as evidence that genetic differences play a bigger role in shaping attitudes as adulthood progresses (Eaves et al. 1997).

We suggest that those genetic factors might include Gendered Fitness Interests. Monozygotic twins are entirely related (i.e. *r* = 1). Moreover, they have identical GFI because they, and their descendent kin, affect one another’s GFI identically. Dizygotic twins, however, diverge in their GFI as soon as one has their first child because that child is related by r=0.5 to the parent, but only r=0.25 to the parent’s twin. Twin registries, coupled with the extensive expertise in variance partitioning among quantitative geneticists, may prove a productive place to begin critical tests of our proposal. One considerable drawback to this idea, however, is the fact that it requires some way for monozygotic twins to identify their genetic unity and thus value their twin as they do their own self. While thinking about inclusive fitness often leads down this path, there is no good basis for expecting that monozygotic twins act in this way.

More generally, our theory of GFI may bear on the question of genetic factors shaping social and political attitudes. Despite strong opinions that such attitudes are acquired environmentally, particularly due to socialization (Miller and Glass 1989, Downey et al. 1994, Eagly et al. 2004, Eagly and Diekman 2006), quantitative genetic analysis, notably from twin studies, has demonstrated substantial additive genetic variance in political and social attitudes (Martin et al. 1986, Eaves et al. 1989, Hatemi et al. 2010). Our theory of GFI adds a hitherto unknown genetic cause of familial resemblance that may, in part, explain the high heritabilities of social and political attitudes that entail a gendered component.

Gendered Fitness Interests are explicitly genetic interests, with relatives exerting an influence on a focal individual’s gendered attitudes in proportion to their genetic relatedness. Relatives that share numbers of mostly male kin (i.e., high *GFI)*, or other sets of relatives that share numbers of mostly female kin (i.e., low *GFI)* will come to resemble one another in GFI more than equally related sets of relatives that share a mix of male and female kin (i.e. *GFI*~0.5). GFI may thus have the effect of amplifying familial resemblance in gendered attitudes more than it would be likely to do for familial resemblance in non-gendered traits such as, say, personality traits, food preferences, or intelligence. If so, GFI may inflate heritability estimates in a way that has, to our knowledge, not been anticipated by quantitative geneticists.

### Paternity uncertainty

Modeling uncertainty of paternity events (Model 3) resulted in smaller variation about the mean GFI for men, and for older women. The effect of kin on GFI was discounted for each instance of paternity, including the paternity involving an individuals’ sons. Hence the reduced variation if GFI in older women. The effects of paternity in more complex models that trace more types of kin, like Model 4, remain to be assessed. The key insight provided by Model 3 is that uncertain paternity dilutes the effects of kin on GFI, as one would likely intuit.

### Family size and GFI

Sex differences in GFI are not affected by family size, as kin are as likely to cause GFI to go down (female kin) as up (male kin). We expected *a priori* that variance in GFI would be smaller when family sizes are large because large numbers of independent events (i.e. sexes of kin) should reduce stochastic variation. Instead, our Model 4 showed that larger families are associated with greater variation about the mean GFI. This happens because large families include the possibility of large numbers of kin of one sex and few if any of the other. The same kinds of large deviations are less frequent when mean family sizes are smaller.

We were interested to know, before beginning this modeling exercise, whether, after including non-descendent kin, any meaningful variation in GFI would remain. From Model 4 it appears that the self and first-order relatives like offspring and siblings (*r* = 0.5) exert sufficient leverage on GFI to make lower-order relatives less important. This model provides some evidence that if one has only data on these kinds of close relatives, one can still estimate GFI with some accuracy.

### Future opportunities for study

We hope that our preliminary suggestions, despite their simplicity, stimulate further discussion and theoretic development. Whether the ability to hold socio-political attitudes that suit one’s kin can evolve via kin Hamiltonian kin selection remains a challenging question far beyond the scope of this paper. Since Hamilton’s (1964a, b) seminal papers, kin selection has benefitted from more than half a century of intense theoretic development and empirical testing. It has generated powerful, original insights into human behavior, including sibling altruism and attitudes concerning inbreeding among kin that have been upheld by empirical tests (Lieberman et al. 2003, 2007, Antfolk et al. 2018). Should the ideas about GFI that we outline here hold up to direct empirical testing, more refined approaches will be needed to discern the mechanisms and the evolutionary dynamics.

For ideology to form the kind of ‘altruism’ that Hamilton sought to explain via kin selection, an individual’s ideological positions would have to influence relatives’ fitness in a manner that obeyed Hamilton’s Rule. That may be unlikely to occur at the level of a community or society, where the efforts of those with high or low GFI will likely cancel out under most circumstances. At the family level, however, we may find that socio-political beliefs could tip behaviors in favor of one sex or another. We suggest that this may be occurring in our study of Islamic veiling practiced in Tunisian households (Blake et al. 2018).

The benefits of ‘altruism’ can depend on dispersal rates and population viscosity, both of which influence the probability of altruism being directed at kin (Hamilton 1964b). Those same benefits can be undermined, however, by the increased competition among relatives that results from an actor’s altruism (West et al. 2002). Whether a socio-political position could take the shape of Hamiltonian altruism, then, depends not only on the positive effects of that position on relatives’ fitness, but also on the countervailing effects of any increase in competition among relatives. A position that favored one daughter might be undermined, for example, if it intensified competition among a family of sisters (i.e. multiple daughters). This notion would need to be accounted for in any argument that socio-political positions are subject to selective forces.

Dispersal, too, might provide the opportunity to separate the effects of kin via intuited GFI versus other more proximate cognitive and social mechanisms involved in knowing and interacting with one’s kin, affines, and step kin etcetera. For example, human kin detection and subsequent behavior toward kin is shaped by cues of relatedness in the form of the time siblings coreside, and associations between infants and mothers (Lieberman et al. 2007). Data that not only tracks the birth and death of relatives, affines, step kin, and other family members, as well as the time spent with them, distance they live away, or simply emotional closeness, might help to disentangle the mechanisms underpinning why the sex of one’s children—and possibly other kin—affects attitudes.

It is possible, perhaps likely, that people are not intuiting genetic relatedness at all, but rather the blend of male and female relatives around them. If this is the case then we would expect step and adoptive siblings, parents, etcetera to influence a person’s attitudes, or at least to do so in proportion to how emotionally close they are to a person. People might simply intuit what is in the interests of their closest geographic or emotional allies, and their attitudes shift to reflect those interests. That would not rule out the possibility that GFI influences socio-political attitudes in an adaptive way. The mechanism of kin recognition need not be perfect in order for kin-biased behavior to evolve (Hamilton 1964a, b, Dawkins 1976, Lieberman et al. 2007, Gardner et al. 2011).

The possibility of kin selection via gendered fitness interests shaping the positions people take on socio-political and ideological issues represents a hypothesis that we believe is worth testing, particularly if big data and greater computing power permit powerful tests of what would probably only constitute small effects. If upheld, Gendered Fitness Interests might go some way to explaining not only why social and political positions are so strongly held, so variable among people, and so likely to shift with age, but also why some people project their ideological positions so avidly. It might also undermine the idea that the interests of individual women and men are perennially, and necessarily at odds. Most people’s Gendered Fitness Interests will be close to equity due to large numbers of both female and male kin. Many individuals, however, with have GFI that are aligned more with the opposite sex than with their own. Both of these observations challenge the idea of biological sex as a fixed and immutable element of individual identity.

## Acknowledgment

This research was funded by Australian Research Council grant DP160100459. We thank Barnaby Dixson, Pauline Grosjean and Michael Jennions for discussions during the development of these ideas.

## Notes

### Competing Interest Statement

The authors have declared no competing interest.

### Summary of Updates

This is a revision before final acceptance.

## References

Alexander, R. D. 1982. Biology and the moral paradoxes. Journal of Social and Biological Structures 5:389–395.

Alexander, R. D. 1987. The Biology of Moral Systems. A. de Gruyter, Hawthorne, N.Y.

Alford, J. R., P. K. Hatemi, J. R. Hibbing, N. G. Martin, and L. J. Eaves. 2011. The Politics of Mate Choice. The Journal of Politics 73:362–379.

Alvard, M. S. 2003. Kinship, lineage, and an evolutionary perspective on cooperative hunting groups in Indonesia. Hum Nat 14:129–163.

Antfolk, J., D. Lieberman, C. Harju, A. Albrecht, A. Mokros, and P. Santtila. 2018. Opposition to Inbreeding Between Close Kin Reflects Inclusive Fitness Costs. Frontiers in Psychology 9.

Bell, E., C. Kandler, and R. Riemann. 2018. Genetic and environmental influences on sociopolitical attitudes: Addressing some gaps in the new paradigm. Politics and the Life Sciences 37:236–249.

Berenbaum, S. A., J. E. O. Blakemore, and A. M. Beltz. 2011. A Role for Biology in Gender-Related Behavior. Sex Roles 64:804–825.

Betzig, L., and L. H. Lombardo. 1992. Whos Pro-Choice and Why. Ethology and Sociobiology 13:49–71.

Blake, K. R., M. Fourati, and R. C. Brooks. 2018. Who suppresses female sexuality? An examination of support for Islamic veiling in a secular Muslim democracy as a function of sex and offspring sex. Evolution and Human Behavior 39:632–638.

Borgerhoff Mulder, M., and K. L. Rauch. 2009. Sexual Conflict in Humans: Variations and Solutions. Evolutionary Anthropology 18:201–214.

Bouchard, T. J. 2009. Authoritarianism, religiousness, and conservatics: Is ‘obedience to authority’ the explanation for their clustering, universality, and evolution? Pages 165–180 in E. Voland and W. Schiefenhövel, editors. The Biological Evolution of Religious Mind and Behaviour. Springer, Berlin.

Calvo-Salguero, A., J. M. A. Garcia-Martinez, and A. Monteoliva. 2008. Differences between and within genders in gender role orientation according to age and level of education. Sex Roles 58:535–548.

Cornelis, I., A. Van Hiel, A. Roets, and M. Kossowska. 2009. Age Differences in Conservatism: Evidence on the Mediating Effects of Personality and Cognitive Style. Journal of Personality 77:51–88.

Cronqvist, H., and F. Yu. 2017. Shaped by their daughters: Executives, female socialization, and corporate social responsibility. Journal of Financial Economics 126:543–562.

Dasgupta, A., L. Ha, S. Jonnalagadda, S. Schmeiser, and H. Youngerman. 2018. The daughter effect: do CEOs with daughters hire more women to their board? Applied Economics Letters 25:891–894.

Dawkins, R. 1976. The Selfish Gene. Oxford University Press, Oxford

DeVaus, D., and I. McAllister. 1989. The Changing Politics of Women - Gender and Political Alignment in 11 Nations. European Journal of Political Research 17:241–262.

Diekman, A. B., and M. C. Schneider. 2010. A Social Role Theory Perspective on Gender Gaps in Political Attitudes. Psychology of Women Quarterly 34:486–497.

Donnelly, K., J. M. Twenge, M. A. Clark, S. K. Shaikh, A. Beiler-May, and N. T. Carter. 2016. Attitudes Toward Women’s Work and Family Roles in the United States, 1976-2013. Psychology of Women Quarterly 40:41–54.

Downey, D. B., P. B. Jackson, and B. Powell. 1994. Sons Versus Daughters - Sex Composition of Children and Maternal Views on Socialization. Sociological Quarterly 35:33–50.

Eagly, A. H., and A. B. Diekman. 2006. Examining gender gaps in sociopolitical attitudes: It’s not Mars and Venus. Feminism & Psychology 16:26–34.

Eagly, A. H., A. B. Diekman, M. C. Johannesen-Schmidt, and A. M. Koenig. 2004. Gender gaps in sociopolitical attitudes: A social psychological analysis. Journal of Personality and Social Psychology 87:796–816.

Eagly, A. H., A. B. Diekman, M. C. Schneider, and P. Kulesa. 2003. Experimental Tests of an Attitudinal Theory of the Gender Gap in Voting. 29:1245–1258.

Eaves, L. J., H. J. Eysenck, and N. G. Martin. 1989. Genes, Culture, and Personality: An Empirical Approach. Academic Press, San Diego.

Eaves, L. J., N. Martin, A. C. Heath, R. Schieken, J. Meyer, J. Silberg, M. Neale, and L. Corey. 1997. Age Changes in the Causes of Individual Differences in Conservatism. Behavior Genetics 27:121–124.

Ekehammar, B., and J. Sidanius. 1982. Sex-Differences in Socio-Political Attitudes - a Replication and Extension. British Journal of Social Psychology 21:249–257.

Eysenck, H. J. 1975. Structure of Social-Attitudes. British Journal of Social and Clinical Psychology 14:323–331.

Feather, N. T. 1977. Generational and Sex-Differences in Conservatism. Australian Psychologist 12:76–82.

Fisher, R. A. 1930. The Genetical Theory of Natural Selection. Oxford University Press., Oxford.

Gardner, A., S. A. West, and G. Wild. 2011. The genetical theory of kin selection. Journal of Evolutionary Biology 24:1020–1043.

Gaulin, S. J. C., D. H. McBurney, and S. L. BrakemanWartell. 1997. Matrilateral biases in the investment of aunts and uncles - A consequence and measure of paternity uncertainty. Human Nature-an Interdisciplinary Biosocial Perspective 8:139–151.

Gompers, P. A., and S. Q. Wang. 2017. And the children shall lead: gender diversity and performance in venture capital. Harvard Business School.

Hamilton, W. D. 1964a. Genetical Evolution of Social Behaviour 2. Journal of Theoretical Biology 7:17–34.

Hamilton, W. D. 1964b. Genetical Evolution of Social Behaviour I. Journal of Theoretical Biology 7:1–16.

Hatemi, P. K., J. R. Hibbing, S. E. Medland, M. C. Keller, J. R. Alford, K. B. Smith, N. G. Martin, and L. J. Eaves. 2010. Not by Twins Alone: Using the Extended Family Design to Investigate Genetic Influence on Political Beliefs. American Journal of Political Science 54:798–814.

Huber, G. A., and N. Malhotra. 2017. Political Homophily in Social Relationships: Evidence from Online Dating Behavior. The Journal of Politics 79:269–283.

Hyde, J. S. 2005. The gender similarities hypothesis. American Psychologist 60:581–592.

Jeon, J., and D. M. Buss. 2007. Altruism towards cousins. Proceedings of the Royal Society B-Biological Sciences 274:1181–1187.

Judge, T. A., and B. A. Livingston. 2008. Is the gap more than gender? A longitudinal analysis of gender, gender role orientation, and earnings. J Appl Psychol 93:994–1012.

Lieberman, D., J. Tooby, and L. Cosmides. 2003. Does morality have a biological basis? An empirical test of the factors governing moral sentiments relating to incest. Proc Biol Sci 270:819–826.

Lieberman, D., J. Tooby, and L. Cosmides. 2007. The architecture of human kin detection. Nature 445:727–731.

Lizotte, M. K. 2015. The Abortion Attitudes Paradox: Model Specification and Gender Differences. Journal of Women Politics & Policy 36:22–42.

Lizotte, M. K. 2016. Investigating women’s greater support of the Affordable Care Act. The Social Science Journal 53:209–217.

Lizotte, M. K. 2017. Gender, Partisanship, and Issue Gaps. Analyses of Social Issues and Public Policy 17:379–405.

Lundberg, S. 2005. Sons, Daughters, and Parental Behaviour. Oxford Review of Economic Policy 21:340–356.

Manza, J., and C. Brooks. 1998. The Gender Gap in U.S. Presidential Elections: When? Why? Implications? 103:1235–1266.

Martin, N. G., L. J. Eaves, A. C. Heath, R. Jardine, L. M. Feingold, and H. J. Eysenck. 1986. Transmission of social attitudes. Proceedings of the National Academy of Sciences 83:4364–4368.

Miller, R. B., and J. Glass. 1989. Parent-Child Attitude Similarity across the Life Course. Journal of Marriage and Family 51:991–997.

Oswald, A. J., and N. Powdthavee. 2010. Daughters and Left-Wing Voting. Review of Economics and Statistics 92:213–227.

Perry, G., and M. Daly. 2017. A model explaining the matrilateral bias in alloparental investment. Proceedings of the National Academy of Sciences of the United States of America 114:9290–9295.

Pinker, S. 2002. The Blank Slate: The Modern Denial of Human Nature. Viking, New York.

Pogrebna, G., A. J. Oswald, and D. Haig. 2018. Female babies and risk-aversion: Causal evidence from hospital wards. Journal of Health Economics 58:10–17.

Pratto, F., J. Sidanius, L. M. Stallworth, and B. F. Malle. 1994. Social-Dominance Orientation - a Personality Variable Predicting Social and Political-Attitudes. Journal of Personality and Social Psychology 67:741–763.

Prokos, A. H., C. L. Baird, and J. R. Keene. 2010. Attitudes about Affirmative Action for Women: The Role of Children in Shaping Parents’ Interests. Sex Roles 62:347–360.

Rice, T. W., and D. L. Coates. 1995. Gender-Role Attitudes in the Southern United-States. Gender & Society 9:744–756.

Schmitt, D. P. 2015. The evolution of culturally-variable sex differences: men and women are not always different, but when they are…it appears not to result from patriarchy or sex role socialisation. in T. K. Shackelford and R. D. Hansen, editors. The Evolution of Sexuality. Springer.

Schmitt, D. P., A. E. Long, A. McPhearson, K. O’Brien, B. Remmert, and S. H. Shah. 2017. Personality and gender differences in global perspective. International Journal of Psychology 52:45–56.

Shackelford, T. K., and A. T. Goetz. 2012. The Oxford Handbook of Sexual Conflict in Humans. Oxford UP, New York, NY.

Shafer, E. F., and N. Malhotra. 2011. The Effect of a Child’s Sex on Support for Traditional Gender Roles. Social Forces 90:209–222.

Sidanius, J., and B. Ekehammar. 1980. Sex-Related Differences in Sociopolitical Ideology. Scandinavian Journal of Psychology 21:17–26.

Sidanius, J., F. Pratto, and L. Bobo. 1994. Social-Dominance Orientation and the Political Psychology of Gender - a Case of Invariance. Journal of Personality and Social Psychology 67:998–1011.

Stearns, S. C. 1992. The Evolution of Life Histories. Oxford University Press, Oxford.

Twenge, J. M. 1997. Changes in masculine and feminine traits over time: A meta-analysis. Sex Roles 36:305–325.

Warner, R. L. 1991. Does the Sex of Your Children Matter - Support for Feminism among Women and Men in the United-States and Canada. Journal of Marriage and the Family 53:1051–1056.

Warner, R. L., and B. S. Steel. 1999. Child rearing as a mechanism for social change - The relationship of child gender to parents’ commitment to gender equity. Gender & Society 13:503–517.

Washington, E. L. 2008. Female socialization: How daughters affect their legislator fathers’ voting on women’s issues. American Economic Review 98:311–332.

West, S. A., I. Pen, and A. S. Griffin. 2002. Conflict and cooperation - Cooperation and competition between relatives. Science 296:72–75.

Wood, W., and A. H. Eagly. 2012. Biosocial Construction of Sex Differences and Similarities in Behavior. Advances in Experimental Social Psychology, Vol 46 46:55–123.

